# Occurrence, functionality, and abundance of the *TERT* promoter mutations

**DOI:** 10.1101/2021.05.03.442397

**Authors:** Sivaramakrishna Rachakonda, Jörg D. Hoheisel, Rajiv Kumar

**Author notes:** **Correspondence to:** Rajiv Kumar, Division of Functional Genome Analysis, German Cancer Research Center, Im Neuenheimer Feld 580, 69120 Heidelberg, Germany.

## Abstract

Telomere shortening at chromosomal ends due to the constraints of the DNA replication process acts as a tumor suppressor by restricting the replicative potential in primary cells. Cancers evade that limitation primarily through the reactivation of telomerase via different mechanisms. Mutations within the promoter of the telomerase reverse transcriptase (*TERT*) gene represent a definite mechanism for the ribonucleic enzyme regeneration predominantly in cancers that arise from tissues with low rates of self-renewal. The promoter mutations cause a moderate increase in *TERT* transcription and consequent telomerase upregulation to the levels sufficient to delay replicative senescence but not prevent bulk telomere shortening and genomic instability. Since the discovery, a staggering number of studies and publications have resolved the discrete aspects, effects, and clinical relevance of the *TERT* promoter mutations. The promoter mutations link transcription of *TERT* with oncogenic pathways, associate with markers of poor outcome, and define patients with reduced survivals in several cancers. In this review, we discuss the occurrence and impact of the promoter mutations and highlight the mechanism of *TERT* activation. We further deliberate on the foundational question of the abundance of the *TERT* promoter mutations and a general dearth of functional mutations within noncoding sequences, as evident from pan-cancer analysis of the whole-genomes. We posit that the favorable genomic constellation within the *TERT* promoter may be less than a common occurrence in other noncoding functional elements. The evolutionary constraints limit the functional fraction within the human genome, hence the lack of abundant mutations outside the coding sequences.

## Introduction

Telomeres, which mask natural chromosomal ends from DNA damage repair machinery to maintain genomic integrity, comprise hexanucleotide repeats that range 10-15 kb in length^1, 2^. The mainly double-stranded telomere repeat sequences end in a single-stranded G-rich 150-200 nucleotide long 3’ tail^3, 4^. Intrinsically unstable telomeres represent fragile sites stabilized by the shelterin complex components, DNA binding TRF1, TRF2, POT1, and adaptors TIN2, TPP1, and RAP1^5, 6^. Shelterin complex shields telomeres through mechanisms that differ significantly in pluripotent and somatic tissues^2, 7–10^. Telomeres require a minimum number of tandem repeats for sufficient associated proteins to form a dynamic protective nucleoprotein structure to overcome the end-protection problem^3, 11, 12^.

An intrinsic limitation of the DNA replication process manifests itself during the synthesis phase of a cell cycle due to unfilled gaps at the chromosomal 5′-terminals following the removal of RNA primers^5, 13^. Besides, the inability of the DNA Pol α-primase complex to initiate the replication from the very end of linear DNA on the lagging strands contributes to the loss of ~70-250 nucleotides per cell division, presaging the gradual telomere attrition^14, 15^. That process consequentially generates the G-rich single-strand overhangs with the length proportional to telomere shortening^16^. The single-stranded overhangs are critical for telomere protection through T-loop formation and elongation via telomerase recruitment that involves TRF2, 5’-3’ exonuclease Apollo, and RTEL1 helicase^17–19^. The erosion of chromosomal ends through successive mitoses constraints the replicative potential of primary cells that tumors adapt to evade through telomere stabilization^20–23^. The age-dependent progressive attrition of chromosomal ends, accentuated through genetic defects or exogenous factors, causes telomere dysfunctions that drive the hallmarks of aging, which include cellular senescence, stem cell exhaustion, genomic instability, mitochondrial dysfunction, epigenetic dysregulation, loss of proteostasis, altered nutrition sensing, and inflammation^24–27^. Inherited mutations in genes that encode proteins involved in telomere structure, repair, replication, and preservation of equilibrium result in debilitating syndromes collectively termed telomeropathies^28, 29^.

Telomere maintenance and stabilization entails a complex and controlled process involving several components functional in critical pathways^30^. The ribonucleic protein telomerase, comprised of intricately interlocked catalytic reverse transcriptase subunit (TERT) and an RNA component (TERC) along with auxiliary elements including histone H2-H2B dimer, counteracts telomere shortening to overcome the ‘end replication problem’ and maintain genomic integrity in pluripotent stem cells, early embryonic tissues, and cells that undergo divisions as a physiological requirement^1, 31–34^. With a limited amount of both the enzyme and substrates, telomerase extends telomeres in the late S-phase through stringent mechanisms “where any perturbation becomes causal for different telomere related diseases, insufficiency leading to stem cell and tissue failure syndromes and too much to cancer predisposition”^35–41^. The recruitment and processivity of telomerase on telomeres are assisted by the shelterin components and terminated by the heterotrimeric CTC1-STN1-TEN1 (CST) complex, followed by a C-strand fill-in the engagement of DNA polymerase α-primase ^37, 42–45^. Most human somatic tissues and adult stem cells do not express sufficient telomerase to maintain telomere length infinitely due to repression of *TERT* upon differentiation in a histone deacetylase-dependent manner and through alternative splicing of TERT expression^27, 46, 47^. Splicing out of exon 2 of *TERT* leads to mRNA decay in differentiated cells, and its retention promotes telomerase accumulation in pluripotent cells^47^. The age-dependent telomere attrition leading to induction of DNA damage response acts as a tumor suppressor mechanism through upregulation of checkpoint inhibitors^3, 48–50^.

The unlimited replicative potential of tumor cells through stabilized telomeres constitutes one of the cancer hallmarks^15, 22, 51^. The stabilization of critically short telomeres through telomerase upregulation or less typically via homologous recombination-based alternate lengthening of telomeres allows tumor cells to escape replicative senescence or tide over the replicative crisis to continued cell divisions through the stages of cancer progression^2, 52^. The detection of recurrent genetic alterations associated with repeat preservation at chromosomal ends through large-scale pan-cancer whole-genome studies has highlighted the importance of the telomere maintenance mechanism in cancers^53, 54^. Telomerase reactivation in tumors occurs through several mechanisms, including amplification, rearrangements, viral integrations, and promoter methylation at the locus leading to increased *TERT* transcription^55–60^. The discovery of the mutations within the core promoter of the *TERT* gene, described as a milestone in telomere biology, afforded a definite mechanism of telomerase upregulation^61–63^. Those noncoding mutations, abundant mainly in cancers that arise from tissues with low rates of self-renewal, cause a moderate increase in *TERT* transcription to rejuvenate telomerase to the levels that delay replicative senescence^61, 62, 64–67^.

This review discusses the occurrence and impact of the promoter mutations and highlights the mechanism through which those alterations upregulate *TERT* and reactivate telomerase. We further deliberate on the foundational question of the abundance of the *TERT* promoter mutations in tumors from tissues with low rates of self-renewal and a general dearth of noncoding mutations in cancers, as evident from pan-cancer analysis of the whole genomes^68^.

## Discovery of the *TERT* promoter mutations

The *TERT* promoter mutations were discovered simultaneously through two different approaches^61, 62^. One group found the somatic *TERT* promoter mutations by analyzing whole-genome data from tumors of unrelated melanoma patients followed by validation in an extended set of tumors^62^. The second group arrived at those mutations prompted by discovering a highly penetrant germline mutation in a large melanoma family through linkage analysis^61^. Both groups identified high-frequency mutually exclusive heterozygous C>T mutations at −124 (1,295,228) and −146 (1,295,250; Figure 1) bp positions from the ATG start site. Both mutations create *de novo* binding sites for ETS transcription factors with the GGAA/T as a general recognition motif and the CCGGAA/T motif for ternary complex factors (TCF)^61, 62^. The investigators that identified the promoter mutations from whole-genome sequencing data also found those alterations in cell lines from various other cancers^62^. The reporter assays with somatic *TERT* promoter mutant constructs displayed a two- to fourfold increase in transcriptional activity over the background of the wild-type sequence^62^.

**Figure 1.**
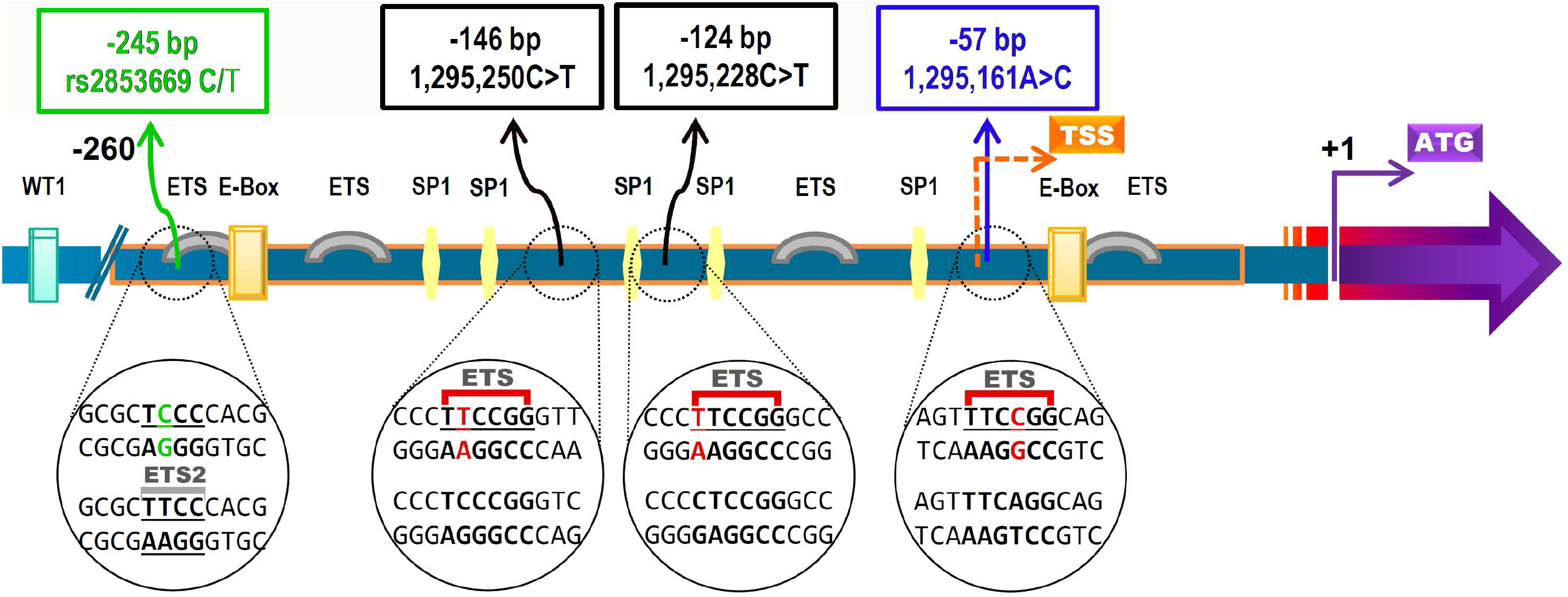
A schematic representation of the core *TERT* promoter. The mutations at −57, −124, and −146 bp positions from the ATG start site of the *TERT* gene create consensus binding motifs for ETS transcription factors. The −124 C>T and −146 C>T somatic mutations are marked in black the germline −57>AC mutation in blue. The variant allele of a common T>C polymorphism represented by rs2853669 at −245 bp disrupts a preexisting non-canonical ETS2 binding site close to an E-box^2^.

The other group initially identified a highly penetrant germline *TERT* promoter mutation through linkage analysis of a large melanoma pedigree with 14 related patients^61^. Targeted sequencing of the disease-linked 2.2 Mb region on chromosome 5p led to the identification of the causal A>C variant at −57 base pairs (bp) position (Chr 5: 1,295,161 hg19 coordinate; Figure 1) from the ATG start site of the *TERT* gene in the affected individuals. The identified single nucleotide transversion (CC<u>T</u>GAA>CC<u>G</u>GAA; +strand) creates a *de novo* binding site for ETS/TCF transcription factors similar to the somatic mutations. The altered base resulted in ~1.5-fold increased luciferase activity in reporter assays over the basal levels with the wild-type sequence^61^.

The mutation carriers within the family developed melanoma with an early age of onset and rapid progression to fatal metastases; two individuals also presented additional malignancies. One person had ovarian cancer at age 27 and melanoma when 30 years old; another individual survived melanoma at age 20, later developed ovarian cancer, renal cell carcinoma, bladder cancer, mammary carcinoma, and finally bronchial carcinoma leading to her death at age 50^61^. The germline TERT promoter mutation detection is cited as a specific example of cancer-predisposing high-penetrant regulatory alteration^69^. A follow-up targeted sequencing initiative involving 675 melanoma families identified the same germline −57 A>C *TERT* promoter mutation in the affected individuals in a single 7-case melanoma family from Leeds^70^. With only two identified families, the germline *TERT* promoter mutation accounts for less than 1% of familial melanoma^48^. Further studies on cancer-prone families also reported disease segregating missense variants in genes that encode shelterin components that included *POT1*, *ACD*, *TRF2*, and *TINF2*^71–74^.

## Distribution of somatic *TERT* promoter mutations

The somatic *TERT* promoter mutations occur in many cancers with an overall frequency of about 27 percent^55^. The frequency of those mutations varies among cancer types and sub-types, with the −124C>T base-change being overwhelmingly dominant in most cancers except for skin neoplasms^2, 57, 75–79^. In melanoma and non-melanoma skin cancers, the −146C>T mutation exceeds the −124C>T in frequency, signifying differences in etiology and distinct mechanisms of mutagenesis^78, 80–85^. The CC>TT tandem mutations at −124/−125 and −138/−139 bp positions from the ATG start site create identical binding motifs, occur specifically in skin cancers, and constitute about 10 percent of the detected *TERT* promoter alterations^61, 80, 86, 87^. The −138/−139 CC>TT mutation also appears infrequently in non-skin cancers as the base-change at −139 bp is reported as a rare C>T polymorphism represented by rs35550267^2, 88^. Other non-frequent somatic mutations within the *TERT* promoter detected in different cancers include the −124C>A and −57C>A, with the latter initially described as the causal familial germline mutation^61, 88^.

The *TERT* promoter mutations predominantly occur in tumors from tissues with low rates of self-renewal due to limiting telomerase levels in the transforming cells^65, 75^. The acquisition of the promoter mutations in melanoma, liposarcoma, hepatocellular carcinoma, urothelial carcinoma, and medulloblastoma due to lack of telomerase becomes advantageous for instantaneous proliferation^65, 75, 78^. For tumors arising from telomerase-positive cells -including cancers of the hematopoietic system, gastrointestinal stromal, lung, ovary, uterine cervix, and prostate- the initial acquisition of *TERT* promoter mutations would unlikely be of any selective advantage^65, 75, 89–94^. That paradigm is supported by the acquisition of the *TERT* promoter mutations as a somatic genetic rescue mechanism in patients with pulmonary fibrosis and aplastic anemia caused by telomerase loss due to germline variants in telomere-related genes^95–97^.

## Impact of *TERT* promoter mutations

Since the discovery, a staggering number of studies and publications have resolved the discrete aspects, effects, and clinical relevance of the *TERT* promoter mutations. Pan-cancer cell lines with the *TERT* promoter mutations exhibit gene-expression characteristics dominated by epithelial to mesenchymal transition and MAPK activation signaling generating distinct tumor environments and intercellular interactions^98^. The promoter alterations associate with cancer stemness through telomerase upregulation, link transcription of the telomerase catalytic subunit with oncogenic pathways, associate with markers of poor outcome, and define patients with reduced survival in different cancers^80, 86, 88, 99, 100^. Those mutations define a switch between adenomas and carcinomas of the liver and are a prerequisite for the rapid growth of tumors in isocitrate dehydrogenase (*IDH*) wild-type adult glioblastomas after years of dormancy^101–103^.

The *TERT* promoter mutations with substantial frequencies occur mainly in specific clinical and phenotypic subtypes. In melanoma, those mutations associate with advanced stages, markers of poor prognosis, increased tumor growth, and, together with *BRAF/NRAS* alterations, define patients with reduced disease-free and melanoma-specific survival^80, 86, 104–107^. A study to measure the effect of individual *TERT* promoter alterations showed that the less frequent −138/−139 CC>TT tandem mutation had the worst impact on disease-free and melanoma-specific survival in stage I and II patients^87^. In different melanocytic neoplasms, the *TERT* promoter mutations serve as ancillary tools for ambiguous histological tumors because of their high specificity for melanoma and absence in melanocytic nevi and melanocytes adjacent to the tumors^64, 108–111^. In bladder cancer, besides being associated with an increased disease recurrence and poor patient outcome, the specificity of the *TERT* promoter mutations distinguishes histologically deceptive cancers from benign mimics^88, 112^. In gliomas, the *TERT* promoter mutations associated with high disease grade, worst patient survival, and other markers follow histological classification into subgroups with distinct disease outcomes^75, 113–119^. An investigation based on editing out of the *TERT* promoter mutations showed that the local injection of adenine base editor fused to Campylobacter jejuni CRISPR-associated protein 9 inhibited the growth of gliomas harboring those alterations^120^. Those mutations cause aggressiveness in meningioma resulting in reduced patient survival and define a highly aggressive subgroup of patients with pleural mesothelioma^121, 122^. The *TERT* promoter mutations in combination with *BRAF* alterations define the most aggressive subtypes among papillary thyroid cancer patients with distance metastases, the highest recurrence, and mortality^123^. In differentiated thyroid cancer, the promoter mutations define an independent prognostic factor after adjusting for risk factors described in the 8^th^ edition of the American Joint Committee on Cancer classification^124^.

The *TERT* promoter mutations tend to occur together with other oncogenic alterations in different cancers with functional consequences. In melanoma and thyroid cancer, the promoter mutations occur more frequently in tumors over the background of *BRAF/NRAS* oncogenic alterations^61, 80, 123^. More than half of the melanoma tumors carrying *BRAF* or *NRAS* alterations acquire *TERT* promoter mutations with synergistic functional consequences^125^. In hepatocellular carcinoma, *TERT* promoter mutations associate positively with *CTNNB1* and *ARID2* mutations as well as *CDKN2A* deletions^126^. *BRAF* mutations render *TERT* expression dependent on MAPK activation in tumors with the promoter mutations. Both alterations linked by ETS1 synergistically promote cancer cell proliferation and immortalization^127, 128^. The RAS-extracellular signal-regulated kinase (ERK) regulates the active chromatin state through physical binding of ERK2 to the mutant *TERT* promoter via displacement of histone deacetylase 1, leading to the recruitment of RNA polymerase II^100, 129^. Activated MAPK also phosphorylates FOS to upregulate GABPB, which in turn binds to the mutant *TERT* promoter^130^. FOS inhibitor T-5224 reportedly suppresses growth in cancer cells and tumor xenografts with the *TERT* promoter mutations by inducing apoptosis, the transcriptional activation of tumor necrosis factor-related apoptosis-inducing ligand receptor 2 (TRAIL-R2), and inactivation of survivin^131^. The presence of *TERT* promoter mutations in cell lines over the background of activated BRAF triggers strong apoptosis-induced cell death upon treatment with MAPK inhibitors that abolishes the growth of *in vivo* tumors harboring both mutations^132^.

A polymorphism rs2853669 at the −245 bp position (Figure 1) from the ATG site within the proximal promoter of the *TERT* gene modulates the effect of the mutations on disease outcome as reported in different cancers^86, 88, 106, 117, 133, 134^. The variant allele disrupts a preexisting non-canonical ETS2 site adjacent to an E-box in the proximal region of the *TERT* promoter^135^. Experiments based on bacterial artificial chromosomes identified that as the upstream of the two ETS sites on the *TERT* promoter. ETV5 binds at those ETS sites through interaction with c-Myc/Max and in conjunction with E-boxes^136^. The core *TERT* promoter consists of 260 base pairs with several transcription factor-binding sites and lacks a TATA box or a similar sequence^136^. Other sequence elements on the *TERT* promoter include five GC-boxes, the binding sites for zinc-finger transcription factor family SP1. The binding of Sp1 and Sp3 to their cognate recognition sites leads to *TERT* transcription only in conjunction with a permissive chromatin environment; whereas, several factors, including p53, down-regulate *TERT* through interaction with Sp1 or other transcription factors^136–138^.

## The functionality of *TERT* promoter mutations

The *TERT* promoter mutations result in increased promoter activity in cells transfected with vectors containing the mutant constructs^61, 62, 88, 127^. The malignant lesions with the promoter mutations from various cancers exhibit a statistically significant enhanced *TERT* transcription and telomerase activity^80, 94, 114, 116, 139–143^. The introduction of the promoter mutations into stem cells prevents the repression of *TERT* at the point of differentiation, and the differentiated cells display telomerase activity comparable to immortal tumor cell lines^65^. Compared to the −57 A>C and −146 C>T mutations, the −124 C>T alteration exerts a maximal effect on *TERT* transcription in tumors and stem cells^65, 114, 144^. In addition, the *TERT* promoter mutations through *de novo* binding sites for ETS transcription factors increase chromatin accessibility, as shown by over-representation of the mutant alleles in ATAC-sequencing assays^145^.

Transcription regulation by ETS factors -a large family with about 27 members reported in humans- involves multi-protein/DNA complexes^136, 146, 147^. In glioblastoma, liver cancer, and bladder cancer cell lines, the obligate multimeric ETS family member GA-binding protein, alpha subunit (GABPA) as a heteromeric complex with GABPB1 binds to the *de novo* consensus sites generated by the −124 C>T and −146 C>T mutations in cooperation with the native sites^148, 149^. The recruitment of GABP transcription factors also mediates a long-range chromatin interaction with sequence elements 300 kb upstream^150^. Studies in multiple cell lines demonstrated an epigenetic switch on the mutant allele to H3K4me2/3, a mark of active chromatin, along with the recruitment of RNA-polymerase II after the binding of GABPA/B1 complex leads to a monoallelic *TERT* expression^149, 151^. The wild-type allele, in contrast, retains H3K27me3, indicating a continued epigenetic silencing^149^. Experiments in glioblastoma cells showed that the disruption of GABPβ1L, a tetramer forming long isomer of GABPB1, decreased *TERT* expression, telomere loss, and cell death in the promoter mutation-dependent manner. In a xenograft model, the decrease in tumor growth in the brains of immunocompromised mice ensued the implantation of human cells with disrupted GABPβ1L^152^. Combined with temozolomide chemotherapy, inducible GABPβ1L knockdown and associated *TERT* reduction reportedly affects DNA damage response leading to reduced growth of intracranial glioblastoma tumors^153^. Other ETS factors that bind those consensus sites on mutant alleles include ETS1 in glioblastoma and ETV5 in GABP-negative cell lines from thyroid cancer^128, 154^. The binding to the site created explicitly by the −146 C>T mutation also involves non-canonical NF-kB signaling with a cooperative binding between p52/RelB and ETS1^155, 156^. The *TERT* promoter mutations also map to the central quadruplex leading to possible alteration of hydrodynamic properties, stability, and local epigenetic modifications^157–159^. Additionally, the promoter mutations reportedly disrupt the binding of non-telomeric TRF2 to the G-quadruplex within the *TERT* promoter leading to telomerase reactivation^160^.

## *TERT* promoter mutations and telomere length

Being causal for increased *TERT* and consequently telomerase levels, it was assumed that the promoter mutations would associate with increased telomere length. Based on the correlation in different human tissues, the telomere length in surrogate tissues like blood reflects the status in tumor-affected organs^161^. Measured in blood cells, individuals from the melanoma family carrying the germline −57 A>C *TERT* promoter mutation had extra-extended telomeres compared to the non-carriers^61, 162^. Similarly, teratomas originating from an injection of hESC with the −124 C>T *TERT* promoter mutation had telomere length comparable to that in undifferentiated cells ^65^. In contrast, telomeres in general, due to the excessive proliferation or other disruptions like shelterin complex dysregulation, are shorter in tumors than in the noncancerous tissues^162–165^. Despite rejuvenated telomerase in most instances, cancers due to functional constraints maintain short telomeres, as the forced elongation results in differentiation and suppression of innate immune-related genes implicated in the maintenance of tumors in an undifferentiated state^166^.

As observed in several cancers, telomeres are invariably shorter in tumors with the acquired *TERT* promoter mutations than those without^55, 114, 126, 134, 162, 167–170^. Earlier it was thought that the acquisition and the context at the telomere crisis during cellular transformation could be a reason for that observation in agreement with the reported reduced telomere contents in tumors with *TERT* modifications^54, 171^. Bypass of replicative senescence and continued telomere attrition in affected cells was postulated to be causal for unstable genomes through cycles of end fusions and breakages with short telomeres causal for initiating cascades leading to chromothripsis and kataegis at the point of crisis^172, 173^. The later evidence pointed to an early occurrence of the *TERT* promoter mutations before biallelic inactivation of checkpoint inhibitors and continued shortening of telomeres in tumors as melanoma^64, 110, 174^.

Based on experiments with isogenic human embryonic stem cells (hESC) experiments, it was proposed that the acquisition of *TERT* promoter mutations “rejuvenates telomerase sufficient to stabilize critically short telomeres to delay replicative senescence but insufficient to prevent bulk telomere shortening”^64^. The continued proliferation in the process elicits genomic instability through an increased number of short telomeres setting the context for decreased telomere length in tumors with those mutations and selection pressure for telomerase upregulation^64^. A further increase in telomerase presumably pulls cells out of telomere crisis through an unknown mechanism with a heavily rearranged genome^3, 64^. The lack of correlation between the *TERT* promoter mutations and telomere length also extends to telomerase levels as the holoenzyme preferentially acts on the shortest telomeres^64, 99^. Transforming cells with relatively long telomeres, limited telomerase, and deleted checkpoints survive critical barriers until the telomerase upregulation. That is explained as an antecedent for the association between increased constitutive telomere length and cancer risk; the evolution of tumors is predicated on the serial accumulation of driver mutations through rapid cell divisions, and the variation in telomere length can affect the proliferative potential of premalignant cells^48, 64, 162, 175–179^.

## The context for the abundance of *TERT* promoter mutations

The discovery of the *TERT* promoter mutations represented two conceptual advancements, genomic alterations driving cancers via transcriptional alteration and a trail of missing mutations not accounted for through only coding sequences^53, 68, 180, 181^. That raised expectations for a surfeit of driver mutations lurking within the functional elements outside the protein-coding sequences in human cancers. The data from the encyclopedia of DNA elements (ENCODE) project stressed the importance of genomic regulatory elements within the human genome and an enhanced understanding of the role of the spatial organization of the genome and cis-acting elements in gene regulation^182, 183^. The mutations within functional noncoding elements, including promoters, enhancers, insulators, long noncoding RNAs, and other regulatory elements, can affect expressions of disparate critical genes^66, 184^. Different initiatives did identify many mutations in the noncoding genome, particularly within the promoter regions of several genes in various cancers, albeit at relatively low frequencies and with less than consequential functional outcomes apart from a few exceptions^66, 68, 185–189^. However, the pan-cancer whole-genome studies based on the extended datasets from 2,658 genomes imply that independent of statistical power, noncoding cis-regulatory driver mutations in known cancer genes other than those within the *TERT* promoter are much less frequent than in the protein-coding sequences^68^. That foundational anomaly merits an attempt for an epistemological understanding.

The relative overrepresentation of the *TERT* promoter mutations may exist due to a vulnerable milieu at the locus where a single base alteration provides motifs for optimal binding of ETS transcription factors to drive up the transcription^190^. Upregulation of mutant allele-specific *TERT* transcription involves spatial architecture where binding of the GABPA/GABPB1 complexes at the *de novo* sites coopts optimally spaced preexisting tandem proximal native ETS sites^148^. Two in-phase proximal tandem native ETS sites within the *TERT* promoter at 30/53 and 25/48 base-pairs from the mutational sites at −124 and −146 bp from the ATG start site (Figure 2A), respectively, are obligatory for facile binding of the GABP heteromeric complex^148, 149^. The spatial arrangement between native and *de novo* ETS sites permits competitive recruitment of the GABP complex through the displacement of initially coopted ELF1/2 at the latter sites^191^. The fortuitous co-occurrence of the ideally spaced tandem native sites near the mutational hotspots on the *TERT* promoter, crucial for that optimal spatial architecture, may be less than a routine occurrence within promoters or other regulatory elements elsewhere in the genome. In the absence of such permissive settings, selecting a single mutation may not be as consequential as in the *TERT* promoter. The differences in relative proximities and conformations between the preexisting sites and those created by different *TERT* promoter alterations could also be factors for the observed intra-mutational differences in transcription, mechanism, and disease outcomes^87, 114, 155, 156^.

**Figure 2.**
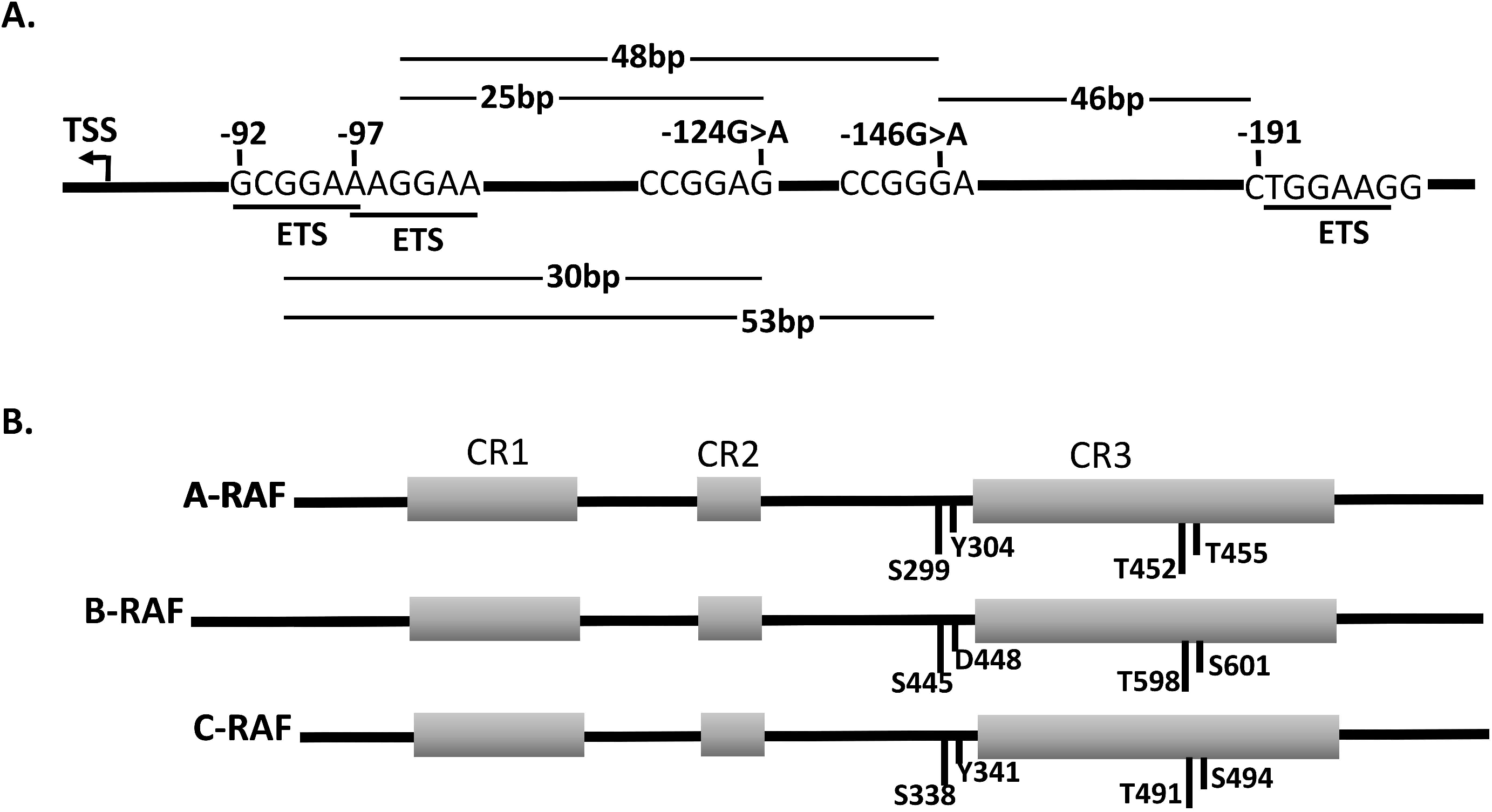
**A.** A schematic representation of the *de novo* binding sites for ETS transcription factors created by the mutations at the −124 and −146 bp sites on the *TERT* promoter. The two preexisting tandem ETS sites 30 and 53 bp from the two mutational sites at −124 and −146 bp, respectively, are required for cooperative binding of the GABP heterotetramer complex at the *de novo* sites^148^. **B.** Structure of the RAF proteins. A single mutation at V600 residue within the third conserved region (CR3) mimics activation of the two adjacent residues is sufficient to activate BRAF. The constitutively phosphorylated S445 residue and the phosphomimetic aspartic acid at the amino acid 448 eliminates the requirement for additional activating alterations needed for maximal activation of ARAF and CRAF paralogs^197^.

The heavy occurrence of a single oncogenic mutation in *BRAF* in melanoma and other cancers and the rareness of alterations at the corresponding residues in *ARAF* and *CRAF* paralogs affords a comparable analogy^192–194^. A single mutation at the V600 residue that mimics activation at two adjacent residues is sufficient to activate *BRAF* ~10-fold over basal levels^195, 196^. Constitutively phosphorylated S445 residue and the 448 residue with the phosphomimetic aspartic acid in *BRAF* (Figure 2B) eliminates the need for additional activating alterations, which would be essential for maximal activation in other RAF isoforms^192, 197^. Similar to the growth advantage accrued from a single nucleotide alteration in *BRAF*, the preexisting native ETS sites within the *TERT* promoter provide a background where the acquisition of a single base change creates an optimally spaced *de novo* ETS site with colossal consequences.

Besides, evolutionary constraints on the fraction of functional noncoding human genome may limit driver mutations outside the coding sequences. Despite the ENCODE biochemical annotations, much of the noncoding sequences other than DNase hypersensitive sites, promoters, and untranslated regions are not optimized for wider pan-mammalian conservation^198, 199^. Instead of a teleological narrative, much of the noncoding human sequences are thought to play nucleotypic or architectural roles without being constricted for specific functionality^199, 200^. For the human population to maintain a constant size through generations, increased fertility must compensate for the reduced mean fitness caused by deleterious mutations. Evolutionary and mutational load constraints arguably limit the functional fraction of the genome to the estimated ~8.2-15%^198, 200–202^. The lack of preexisting favorable constellations in functional elements further restricted by evolutionary constraints could limit the abundant selection of somatic mutations within the noncoding genome. Mutations within the *TERT* promoter fulfill many selection criteria for Darwinian evolutions of tumors that arise in tissues with low rates of self-renewal where upregulation of telomerase delays replicative senescence^203^. The acquisition of the mutations and the consequent *de novo* sites in the human *TERT* promoter is analogous to GABP complex binding sites in somatic cells of rodents, which activate telomerase. The presence or absence of GABPA binding sites in the *TERT* promoter determines whether replicative senescence acts as a tumor suppressor in those species. The *TERT* promoter mutations, as asserted, provide a solution for Peto’s paradox in rodents, based on the observed resistance of large-bodied species to develop cancer compared to small ones^204^.

## Conclusions

The gradual attrition of telomeres due to limited telomerase in most adult cells while quintessential for aging doubles as a natural barrier for cancer development predicated on the serial accumulation of driver mutations. Tumors attain an unlimited replicative potential primarily through telomerase regeneration via increased *TERT* transcription. The *TERT* promoter mutations represent a definite mechanism for telomerase regeneration predominantly in cancers that arise from tissues with low rates of self-renewal. The broadly distributed noncoding mutations have emerged as markers for histological diagnosis, poor disease outcomes in different cancers, and possibly therapeutical targets. The mechanism of the *TERT* activation through the promoter mutations has opened up a range of transcription factors and histone marks that associate with mutant alleles as potential treatment targets. The discovery of the *TERT* promoter mutations, unique in itself, held an unrealized promise of a surfeit of driver mutations in cancers within the noncoding genome. It is plausible that the lack of mutational abundance outside the coding sequences may be due to the lack of fortuitous genomic constellation as that within the *TERT* promoter and evolutionary constraints that limit the functional fraction within the human genome.

## Abbreviations

ACD: Adrenocortical Dysplasia Protein Homolog (unofficial name TPP1)
ARF: A-Raf Proto-Oncogene Serine/Threonine-Protein Kinase
ARID2: AT-Rich Interaction Domain 2
ATAC: Assay for Transposase-Accessible Chromatin
BRAF: B-Raf Proto-Oncogene, Serine/Threonine Kinase
CDKN2A: Cyclin-Dependent Kinase Inhibitor 2A
CRAF: C-Raf Proto-Oncogene, Serine/Threonine Kinase
CST: CTC1-STN1-TEN1 complex
CTNNB1: Catenin Beta 1
ENCODE: Encyclopedia of DNA Elements
ELF: E74 Like ETS Transcription Factor
ERK: Extracellular Signal-Regulated Kinase
ETS: E26 Transformation-specific
ETS1: ETS Proto-Oncogene 1, Transcription Factor
ETS2: ETS Proto-Oncogene 2, Transcription Factor
ETV5: ETS Variant Transcription Factor 5
GABPA: GA Binding Protein Transcription Factor Subunit Alpha
GABPB1: GA Binding Protein Transcription Factor Subunit Beta 1
hESC: Human Embryonic Stem Cells
IDH: Isocitrate Dehydrogenase
MAPK: Mitogen-Activated Protein Kinase
NF-kB: Nuclear Factor Kappa B Subunit
NRAS: Neuroblastoma RAS Viral Oncogene Homolog
POT1: Protection of Telomere
RAP1: Repressor/activator Protein 1
RTEL: Regulator of Telomere Elongation Helicase 1
TERC: Telomerase RNA Component
TERT: Telomerase Reverse Transcriptase
TIN2: TRF1 Interacting Nuclear Factor 2
TRAIL-R2: Tumor Necrosis Factor-related Apoptosis-inducing Ligand Receptor 2
TRF1: Telomeric Repeat Binding Factor 1
TRF2: Telomeric Repeat Binding Factor 2

